# The *queenslandensis* and the *type* form of the dengue fever mosquito (*Aedes aegypti* L.) are genomically indistinguishable

**DOI:** 10.1101/063792

**Authors:** Gordana Rašić, Igor Filipović, Ashley G. Callahan, Darren Stanford, Abigail Chan, Sai Gek Lam-Phua, Huat Cheong Tan, Ary A. Hoffmann

## Abstract

**Background:** The mosquito *Aedes aegypti* (L.) is a major vector of viral diseases like dengue fever, Zika and chikungunya. *Aedes aegypti* exhibits high morphological and behavioral variation, some of which is thought to be of epidemiological significance. Globally distributed domestic *Ae. aegypti* have been traditionally grouped into (i) the very pale variety *queenslandensis* and (ii) the *type* form. Because the two color forms co-occur across most of their range, there is interest in understanding how freely they interbreed. This knowledge is particularly important for control strategies that rely on mating compatibilities between the release and target mosquitoes, such as *Wolbachia* releases and SIT. To answer this question, we analyzed nuclear and mitochondrial genome-wide variation in the co-occurring pale and *type Ae. aegypti* from northern Queensland (Australia) and Singapore.

**Methods/Findings:** We typed 74 individuals at a 1170 bp-long mitochondrial sequence and at 16,569 nuclear SNPs using a customized double-digest RAD sequencing. 11/29 genotyped individuals from Singapore and 11/45 from Queensland were identified as *var. queenslandensis* based on the diagnostic scaling patterns. We found 24 different mitochondrial haplotypes, seven of which were shared between the two forms. Multivariate genetic clustering based on nuclear SNPs corresponded to individuals’ geographic location, not their color. Several family groups consisted of both forms and three *queenslandensis* individuals were *Wolbachia* infected, indicating previous breeding with the *type* form which has been used to introduce *Wolbachia* into *Ae. aegypti* populations.

**Conclusion:** *Aedes aegypti queenslandensis* are genomically indistinguishable from the *type* form, which points to these forms freely interbreeding at least in Australia and Singapore. Based on our findings, it is unlikely that the presence of very pale *Ae. aegypti* will affect the success of *Aedes* control programs based on *Wolbachia*-infected, sterile or RIDL mosquitoes.

**Author Summary:** *Aedes aegypti*, the most important vector of dengue and Zika, greatly varies in body color and behavior. Two domestic forms of this mosquito, the very pale *queenslandensis* and the browner *type*, are often found together in populations around the globe. Knowing how freely they interbreed is important for the control strategies such as releases of *Wolbachia* and sterile males. To answer this question, we used RAD sequencing to genotype samples of both forms collected in Singapore and northern Queensland. We did not find any association between the mitochondrial or nuclear genome-wide variation and color variation in these populations. Rather, “paleness” is likely to be a quantitative trait under some environmental influence. We also detected several *queenslandensis* individuals with the *Wolbachia* infection, indicating free interbreeding with the *type* form which has been used to introduce *Wolbachia* into *Ae. aegypti* populations. Overall, our data show that the very pale *queenslandensis* are not genomically separate, and their presence is unlikely to affect the success of *Aedes* control programs based on *Wolbachia*-infected, sterile or RIDL mosquitoes.

## Introduction

The mosquito *Aedes aegypti* (Linnaeus) is the most important arboviral vector in the tropics and subtropics [1]. Diseases transmitted by *Ae. aegypti*, like dengue fever and Zika, are on the rise [2], and some are reappearing. For instance, chikungunya has returned to the American tropics in 2013, after being absent for nearly 200 years [3]. Yellow fever was nearly eliminated thanks to an effective vaccine, but is now resurging in central and south Africa [4]. Such epidemiological trends highlight the need to persist with vector control efforts, which requires a thorough understanding of vector biology.

Nearly 60 years ago, Mattingly [5] noted that despite a vast body of literature, few mosquitoes have been “*the subject of misconception….in the minds of the general run of entomologists”* like *Aedes aegypti* [5]. The species has a plethora of historical synonyms [6], mainly as a result of having extensive variation in body color and scaling patterns [5][7] which was also thought to correlate with behavioral differences (e.g. [8]). These issues urged Mattingly [5] to revise the taxonomy of *Ae. aegypti* and create a foundation for the modern studies of this disease vector [7].

Mattingly [5] proposed the intraspecific classification of *Ae. aegypti* into three forms.

i. A very dark form that never has pale scales on the first abdominal tergite, avoids biting humans, prefers natural breeding habitats and is confined to sub-Saharan Africa. Mattingly gave this form a subspecies rank, *Ae. aegypti spp. **formosus*** (Walker).
ii. *Ae. aegypti* sensu stricto or the ***type*** form, distinctly paler and browner than *spp. formosus*, with pale scales restricted to the head and the first abdominal tergite. This form prefers to bite humans and to use artificial breeding containers, and is globally distributed.
iii. A very pale form, *Ae aegypti **queenslandensis*** Theobold), with extension of the pale scaling on the thorax, tergites and legs, that co-occurs globally with the *type* form. Mattingly gave this form only a varietal rank (*Ae. aegypty* var. *queenslandensis*).

Because of great variation in color and scaling within and among *Ae. aegypti* populations, McClelland [7] suggested that subdivision into forms seems oversimplistic and should be abandoned unless correlation between genetic and color variation can be demonstrated [7]. His recommendations have been largely disregarded [9] despite the fact that multiple genetic marker systems (allozymes, microsatellites, nuclear and mitochondrial SNPs) have failed to find a clear differentiation between forms and markers [10][11][12].

Recently, Chan et al. [13] suggested that the DNA barcoding technique can be used to distinguish *queenslandensis* individuals from the *type* individuals in Singapore. The sequence divergence of 1.5%-1.9% between the two forms [13], although lower than a commonly adopted threshold of 3% for species delineation in insects [14], suggests that the two forms may not freely interbreed. Historical records indicate that the two forms have co-occurred in Singapore and other parts of south-east Asia and Australia for hundreds of generations [5][8]. In sympatry, genetic isolation can be maintained largely through pre-zygotic isolation mechanisms like incompatibilities in mating behavior [15]. For instance, molecular forms of the malarial mosquito, *Anopheles gambiae*, fly together in mating swarms but rarely hybridize due to flight-tone matching between males and females of the same form [16]. Similar incompatibilities in *Ae. aegypti* would have implications for control strategies that rely on successful mating between the release and target mosquitoes, like *Wolbachia*-based population replacement and suppression [17][18], releases of sterile males [19] or males with a RIDL construct [20].

To explore this further, we analyzed nuclear and mitochondrial genome-wide variation in the co-occurring pale and *type Ae. aegypti* from Singapore and northern Queensland (Australia). The RADseq approach we employed allows for detection of genetic structure and ancestry with power unparalleled by previous genetic studies of the *Ae. aegypti* forms [18]. Any association between genetic structuring (nuclear/mitochondrial) and the mosquito color/scaling would provide support for the hypothesis of restricted interbreeding between the *type* and the *queenslandensis* form, with implications for the implementation of biocontrol programs to suppress diseases transmitted by *Ae. aegypti*.

## Materials and Methods

### Ethics statement

The collection of wild mosquitoes in the study areas does not require specific field ethics approval. The sampling was not conducted on protected land, nor did it involve endangered or protected species. Consent was obtained from residents at each location where collections occurred on private property.

### Sampling and identification

In Singapore, all samples were collected as larvae from the domestic breeding containers at nine locations during the second week of April 2015 (Figure 1, Table 1). These samples were collected during routine inspection by enforcement officers of the National Environment Agency (NEA), Singapore. Larvae were reared to the adult stage under standard laboratory conditions (25° ± 1°C, 80 ± 10% relative humidity and 12 h light/dark cycle). In Townsville (northern Queensland), samples were collected as adults using Biogents Sentinel traps placed at 55 locations in January 2014 (Figure 1, Table 1). Adult mosquitoes were sexed and identified to form based on the key diagnostic color and scaling features, following Mattingly [5] and McClleland [7]. Eleven out of 44 mosquitoes (25%) from Singapore, and seven out of 99 mosquitoes (7%) from Townsville were identified as the *queenslandensis* form (Table 1). Additional four *queenslandensis* individuals collected in Cairns (northern Queensland) in December 2014 were included in the analyses (Table 1).

**Figure 1.**
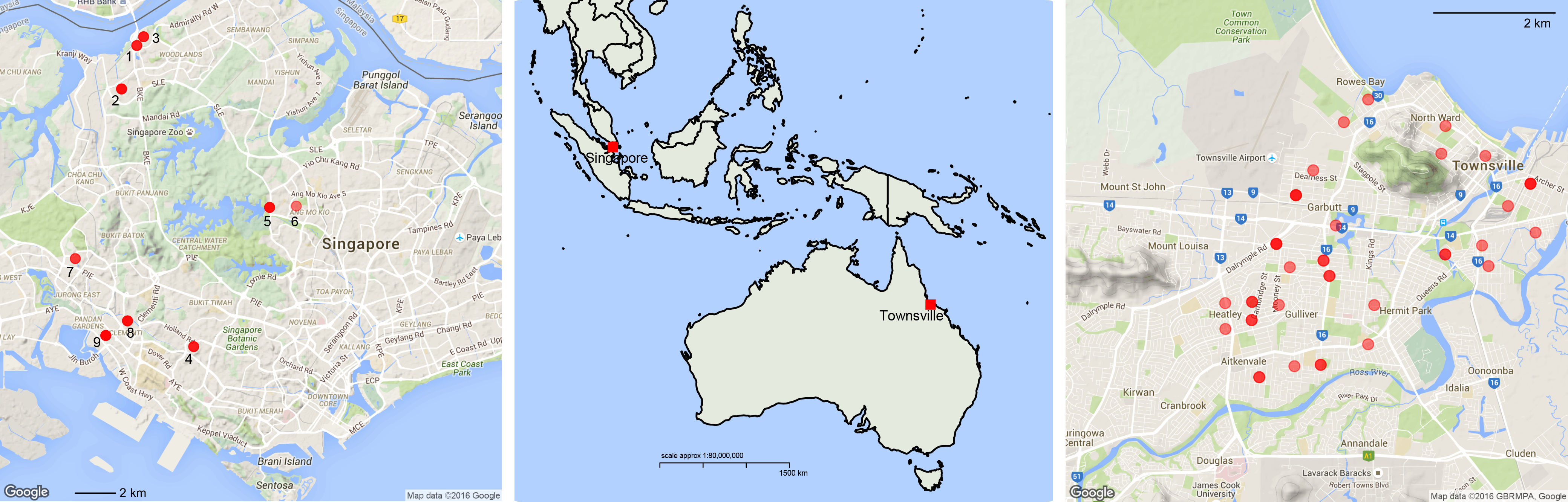
Sampling sites. In Singapore (left), each sampling point represents one breeding container from which larvae were collected. In Townsville, each sampling point represents one BG-Sentinel trap from which adults were collected.

**Table 1.**
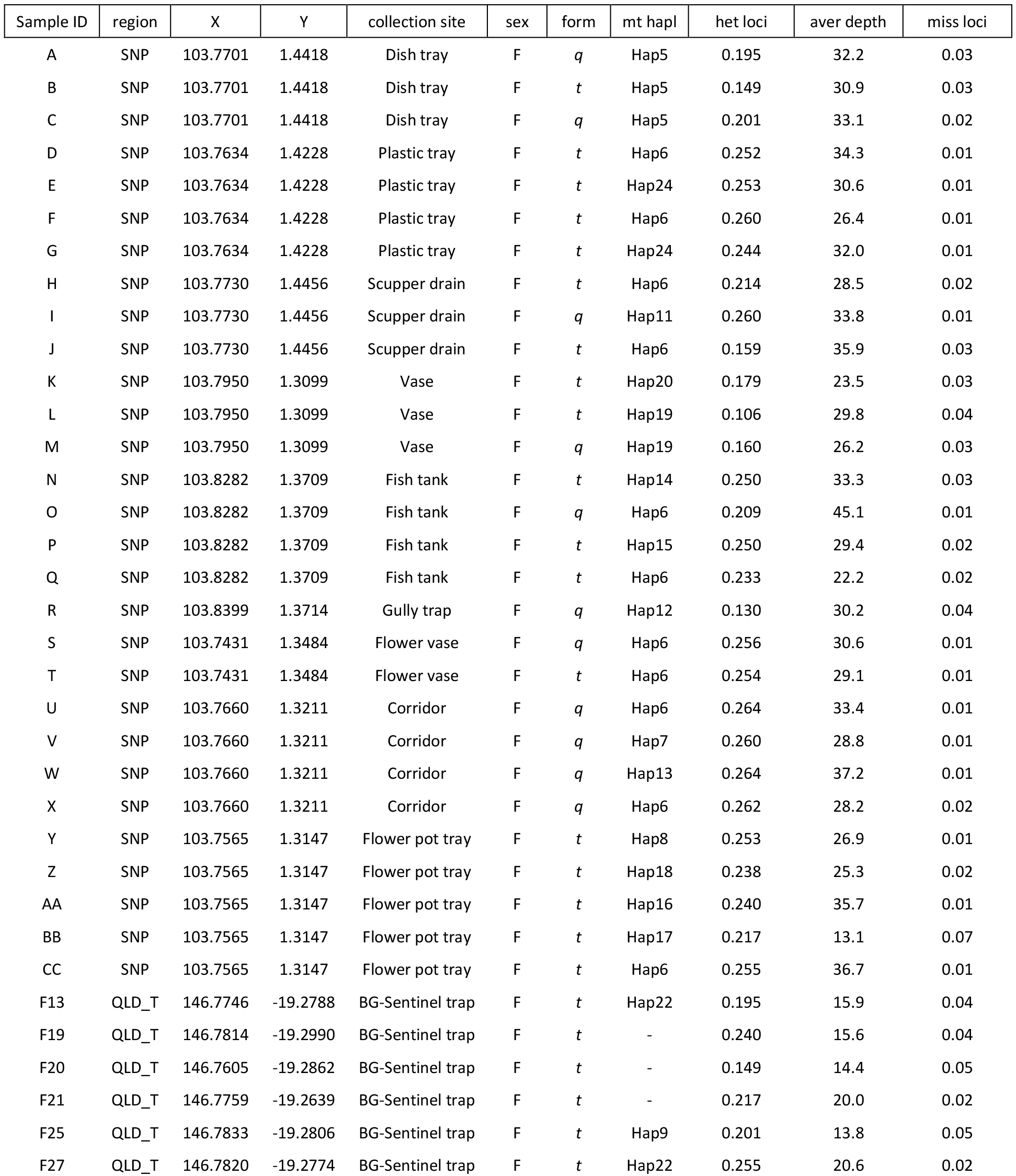
Sample information. Sample ID, region (SNP - Singapore, QLD_T - Townsville, QLD_C - Cairns, Queensland), X, Y (longitude/latitude decimal degrees), collection (method/breeding container), sex (F - female, M - male), form (*t - type, q - queenslandensis* [5][7]), mitochondrial haplotype (mt hapl, Hap1-24), per individual proportion of heterozygous (het) nuclear loci, average (aver) locus depth, and proportion of missing (miss) loci.

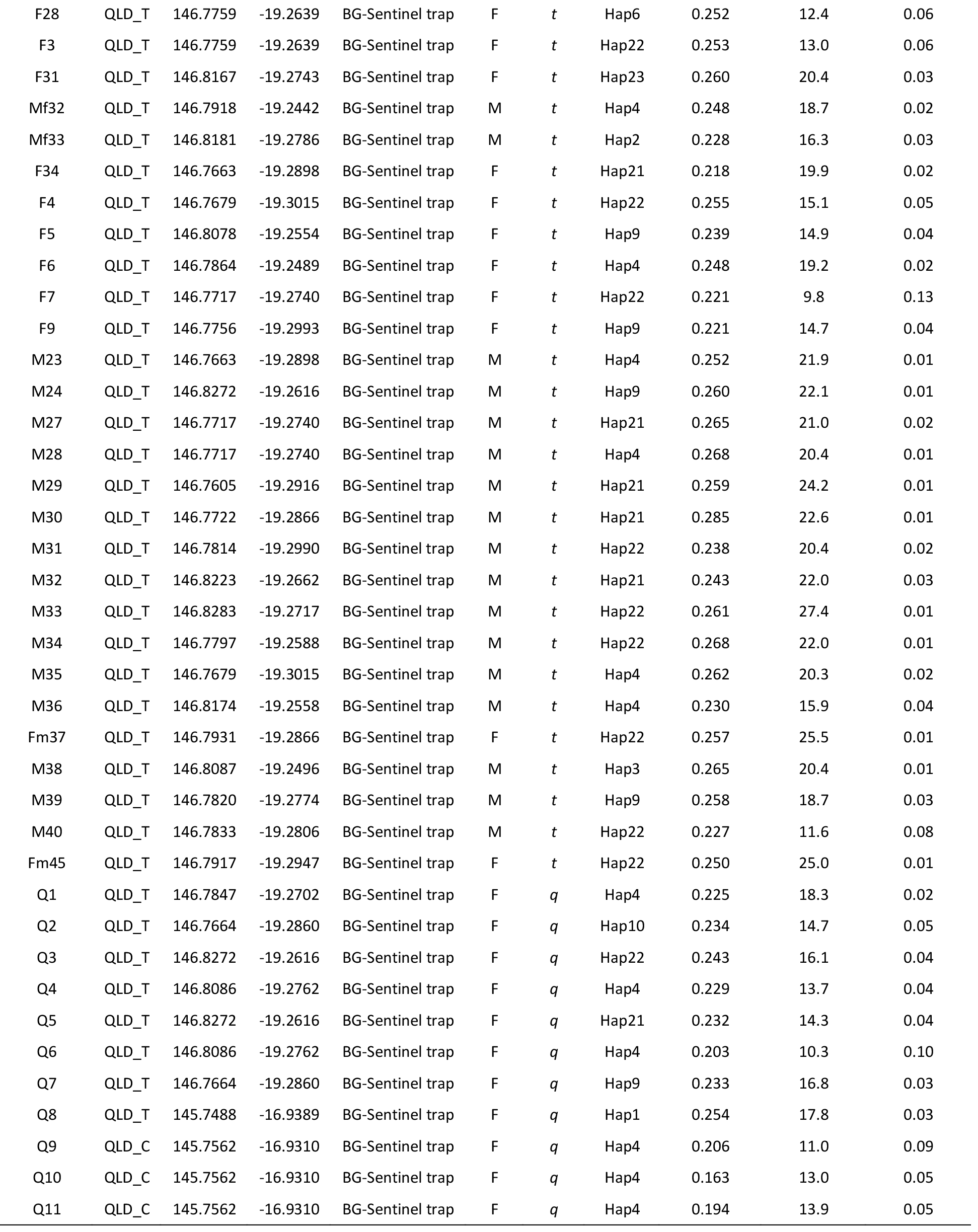

### RADseq genotyping

DNA was extracted from 29 individuals collected in Singapore (18 female *type*, 11 female *queenslandensis*) and 45 individuals from northern Queensland (17 male *type*, 17 female *type*, 11 female *queenslandensis*) (Table 1). Qiagen Blood and Tissue DNA kit (Venlo, Limburg, NL) was used to extract DNA from a whole adult mosquito. 100 ng of DNA from each individual was used to construct the double-digest RAD library following a previously validated protocol [21]. In short, 100 units of the two frequently cutting enzymes (*MluCI* and *NlaIII*, New England Biolabs, Beverly MA, USA) were used to digest 100 ng of DNA during three hours of incubation at 37°C. 100 pM P1 and 300 pM P2 Illumina adapters with customized barcode sequences were ligated to the genomic fragments using 100 units of T4 ligase at 16°C overnight (New England Biolabs, Beverly, MA, USA). Pooled ligations were purified and size selected for fragments 300-450bp in length, using the 2% Pippin Prep cassette (Sage Sciences, Beverly, MA, USA). The final libraries (one for each geographic region) were enriched with 12 PCR cycles with standard Illumina primers and then sequenced in two HiSeq2500 lanes with the 100 bp paired-end chemistry.

Raw fastq sequences were processed within our customized pipeline [21]. First, all reads were trimmed to the same length of 90 bp and removed if the base quality score was below 13 (FASTX Toolkit, http://hannonlab.cshl.edu/fastx_toolkit/index.html). High quality reads were then aligned to the reference mitochondrial genome [22] and the nuclear genome version AaegL1 [23] using the aligner *Bowtie* [24]. Uniquely aligned reads were passed to the *refmap.pl* program that runs the *Stacks* v.1.35 pipeline [25]. In addition to the samples from Singapore, Townsville and Cairns, we included previously sequenced individuals: 15 from Rio de Janeiro (Brazil) [26], 15 from Gordonvale (northern Queensland), and 15 from Ho Chi Minh City (Vietnam) (S1 file). This was done to compare the extent of genetic structuring within and among samples at a regional and global scale. Sexing of the larval samples from Brazil and Vietnam could not be done based on the external morphological features, so we employed a genetic sexing method based on the presence/absence of the male-specific RAD tags [27]. All 119 individuals were included in the creation of the RAD tag catalogues using the default *Stacks* parameters in the maximum likelihood model of SNP and genotype calling. The *populations* module was used to filter the catalogues and export data in the FASTA format (for the mitochondrial variation) and the variant calling format (VCF, for the nuclear variation).

### Analyses of genetic diversity and structure

The mitochondrial haplotype richness within and among groups ( *Ae.aegypti* forms and geographic regions) was calculated using the rarefaction method implemented in the program *HP-rare* [28]. Phylogenetic relationship among mitochondrial haplotypes was estimated with the maximum likelihood approach in the program *RAxML* (GTRM + G, rapid bootstrap heuristic algorithm and thorough ML search) [29]. Haplotypes of three related *Aedes* species, for which the whole mitochondrial genome sequences were available, served as outgroups: *Ae. albopictus* (NCBI: NC_006817.1), *Ae. notoscriptus* (NC_025473.1) [30] and *Ae. vigilax* (KP995260.1) [31]. Haplotype sequence of the *Ae. aegypti* reference line (Liverpool, NC_010241.1) was also included in the analysis.

Parameters of data quality and diversity (RAD tag depth, percentage of missing data, heterozygosity averaged per individual) were compared between females of the two co-occurring forms using independent sample *t*-test. The level of nuclear genetic structuring was estimated using the non-spatial multivariate method DAPC [32] in the *R* package *adegenet* [33]. Rousset’s genetic distance (*â*) and geographic distance between pairs of individuals were calculated in the program *spagedi* [34]. Color distance between pairs of individuals was treated as a binary value: 0 (same color/form) and 1 (different color/form).

## Results & Discussion

### Variation and phylogenetic relationship among mitochondrial haplotypes

From the mitochondrial RAD tag catalogue, we extracted 13 polymorphic tags that were shared between at least 80% of individuals (60/74, Table 1). Tags were distributed across eight different mitochondrial genes (*COXI, Cytb, ATP6, ND1-2, ND4-6;* S2 file). All 13 tags were concatenated into a final 1170 bp long sequence that was treated as a mitochondrial haplotype. We found 24 different haplotypes in samples from Singapore and Townsville. Haplotype richness did not differ between the two forms in either location (Singapore *type* = 5.13, *queenslandensis* = 5.07; Townsville *type* = 4.19, *queenslandensis* = 5.0). Moreover, seven haplotypes were shared between the two forms (Table 1, Figure 2).

**Figure 2.**
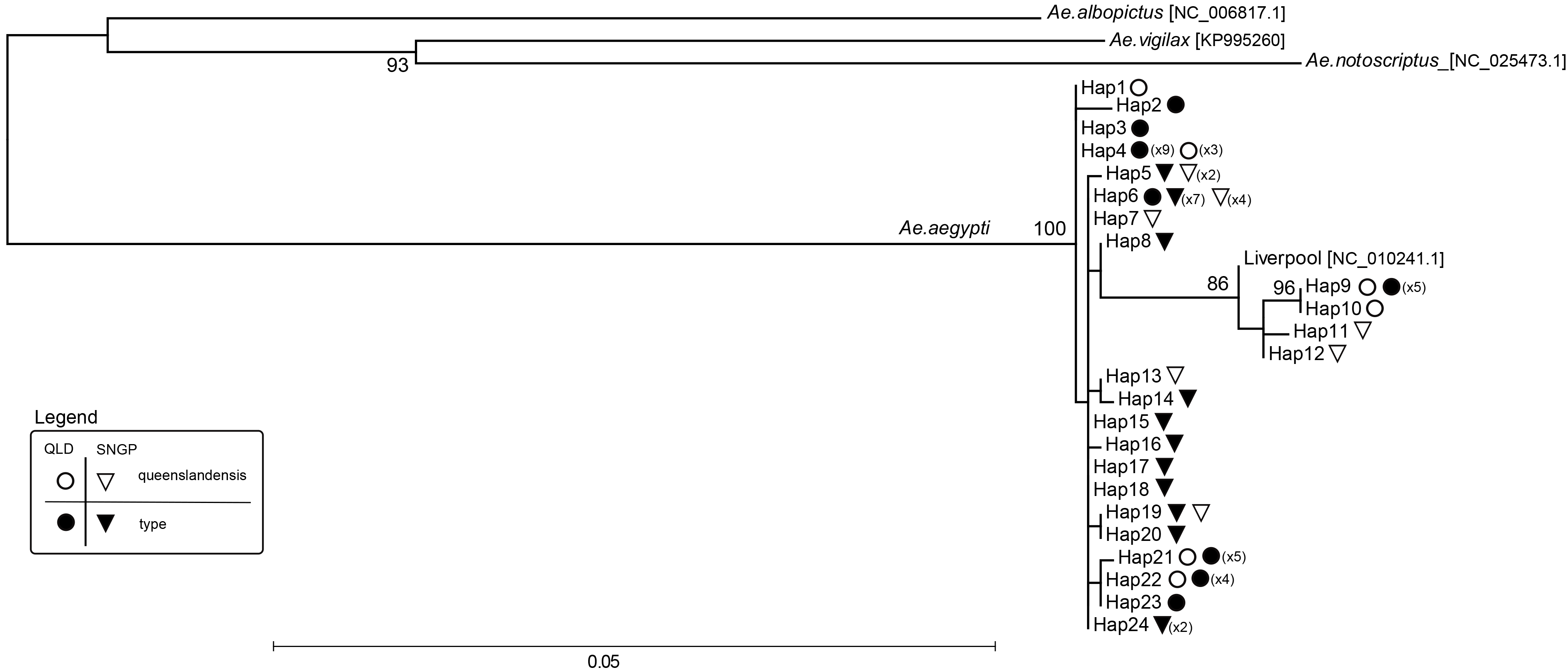
Mitochondrial Maximum likelihood phylogeny. Twenty four different mitochondrial haplotypes (Hap1-24) found in *Aedes aegypti type* and var. *queenslandensis* that co-occur in Singapore and northern Queensland, Australia. Sequences of the three outgroups (*Ae. albopictus, Ae. vigilax, Ae. notoscriptus*) and *Ae. aegypti* Liverpool strain were obtained from the NCBI nucleotide sequence/genome database, with the NCBI accession numbers listed in square brackets. The number of *Ae. aegypti* individuals with a given mitochondrial haplotype is listed in parentheses. A circle designates haplotypes found in Queensland, and a triangle those found in Singapore. Open symbols designate haplotypes found in the *queenslandensis* form, and filled symbols those found in the *type* form.

There were 207 distinctive alignment patterns and 8.17% of undetermined characters in the dataset consisting of 24 haplotypes from Singapore and Queensland, one from the Liverpool strain and three from other *Aedes* species (outgroups). Maximum likelihood phylogeny revealed two highly statistically supported maternal lineages in *Ae. aegypti*: a basal clade (more similar to the outgroups) and a clade arising from it (a derived clade) (Figure 2). Nucleotide distance (*p*-distance) between the two clades ranged from 1.2% to 1.6% (S3 file). Importantly, haplotypes of the two *Ae. aegypti* forms were found in both clades, indicating no association between mitochondrial variation and color variation (Figure 2).

While our results do not support the tentative patterns suggested by Chan et al. [13], they match those from the most comprehensive mitochondrial phylogeny of the African and global *Ae. aegypti* generated to date [35]. Using the ND4 variation, Moore et al. [35] showed that *Ae. aegypti* populations outside Africa represent “mixtures” of mosquitoes from the basal clade and the derived clade, with the basal clade likely originating from West Africa and the derived clade mainly from East Africa. Our analyses of the mitochondrial genome-wide variation revealed the same matrilineage structure in populations from Singapore and northern Queensland (Figure 2). A lack of mitochondrial distinctiveness between the *queenslandensis* and the *type* form is also in line with the findings of More et al. [35], who could not separate the *type* and *formosus* forms into distinct mitochondrial clades despite their assumed subspecies rank.

### Nuclear genetic structuring

We extracted nuclear RAD tags that were shared between at least 80% of individuals in the entire dataset (Singapore, Townsville, Gordonvale, Ho Chi Minh City and Rio de Janeiro). To avoid redundant information from the highly linked markers, we randomly selected one SNP per tag with a minor allele frequency greater than 5%, which gave a total of 16,569 markers for downstream analyses.

Parameters of data quality and diversity did not significantly differ between the co-occurring *queenslandensis* and *type* individuals, including the average: percentage of reads uniquely aligned to the reference genome (Singapore: *t*_df,27_ = 1.46, *p* = 0.15; Townsville: *t*_df,26_ = 0.782, *p* = 0.44), locus depth (Singapore: *t*_df,27_ = 1.66, *p* = 0.11; Townsville: *t*_df,26_ = −1.73, *p* = 0.095), percentage of missing data (Singapore: *t*_df,27_ = −0.67, *p* = 0.51; Townsville: *t*_df,26_ = 0.951, *p* = 0.35), or heterozygosity (Singapore: *t*_df,27_ = 0.46, *p* = 0.65; Townsville: *t*_df,26_ = −2.42, *p* = 0.023) (Table 1, S1 figure).

Discriminant analysis of principal components (DAPC) showed a clear-cut differentiation of mosquitoes based on their geographic origin and not their color. When the entire dataset was considered, *Ae. aegypti* individuals formed genetic clusters that corresponded to their sampling region (i.e. Rio de Janeiro, Ho Chi Minh City, Singapore and northern Queensland) (Figure 3a). The only exceptions were three individuals in Singapore (K,L,M) that formed a distinct genetic group (Figure 3a). They were collected as larvae from the same breeding container, and two were identified as the *type* and one as the *queenslandensis* form (Table 1, Figure 3b). Given their high relatedness (Supplemental file 4) and shared mitochondrial haplotype, as well as high nuclear differentiation from other mosquitoes in the region, it is likely that individuals K, L and M are offspring of the incursion female(s) not local to Australia and Vietnam. These individuals were found near the city port (Figure 1), suggesting a possible route of introduction.

**Figure 3.**
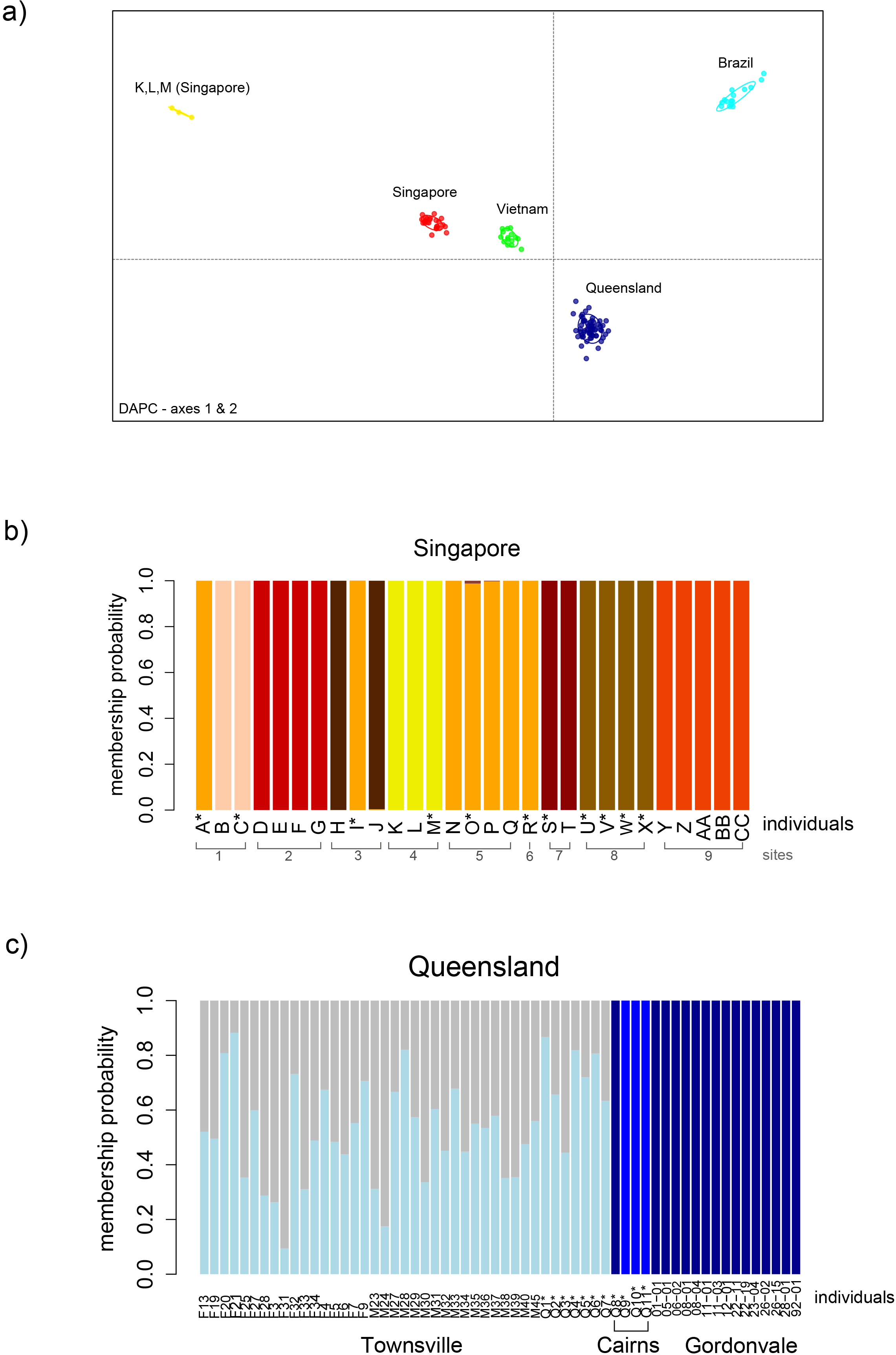
Nuclear genetic structuring (DAPC) Individuals marked with the asterix (*) in their sample ID were identified as *Aedes aegypti queenslandensis* based on the diagnostic scaling patterns [7]. (a) Scatterplot summarizing the individual DAPC scores (axes 1 and 2) in *Aedes aegypti* samples collected in Singapore, Queensland, Ho Chi Minh City (Vietnam) and Rio de Janeiro (Brazil); (b) Individual membership probability to the genetic groups in Singapore; (c) Individual membership probability to the genetic groups in northern Queensland.

Further analysis of genetic structuring within Singapore revealed that family groups were sampled within the breeding containers, some of which had both color forms (Figure 3b). Highly related *queenslandensis* and *type* pairs were found at 4 locations (Figure 3b), including the incursion family group (K,L,M). Most of the related individuals (24/28 pairs), however, had the same color (Figure 4). These results suggest that the color/scaling pattern is likely to represent a quantitative trait under some environmental influence, which is in line with the recent discovery of multiple QTLs associated with the dorsal abdominal scaling pattern in *Ae. aegypti* [36].

**Figure 4.**
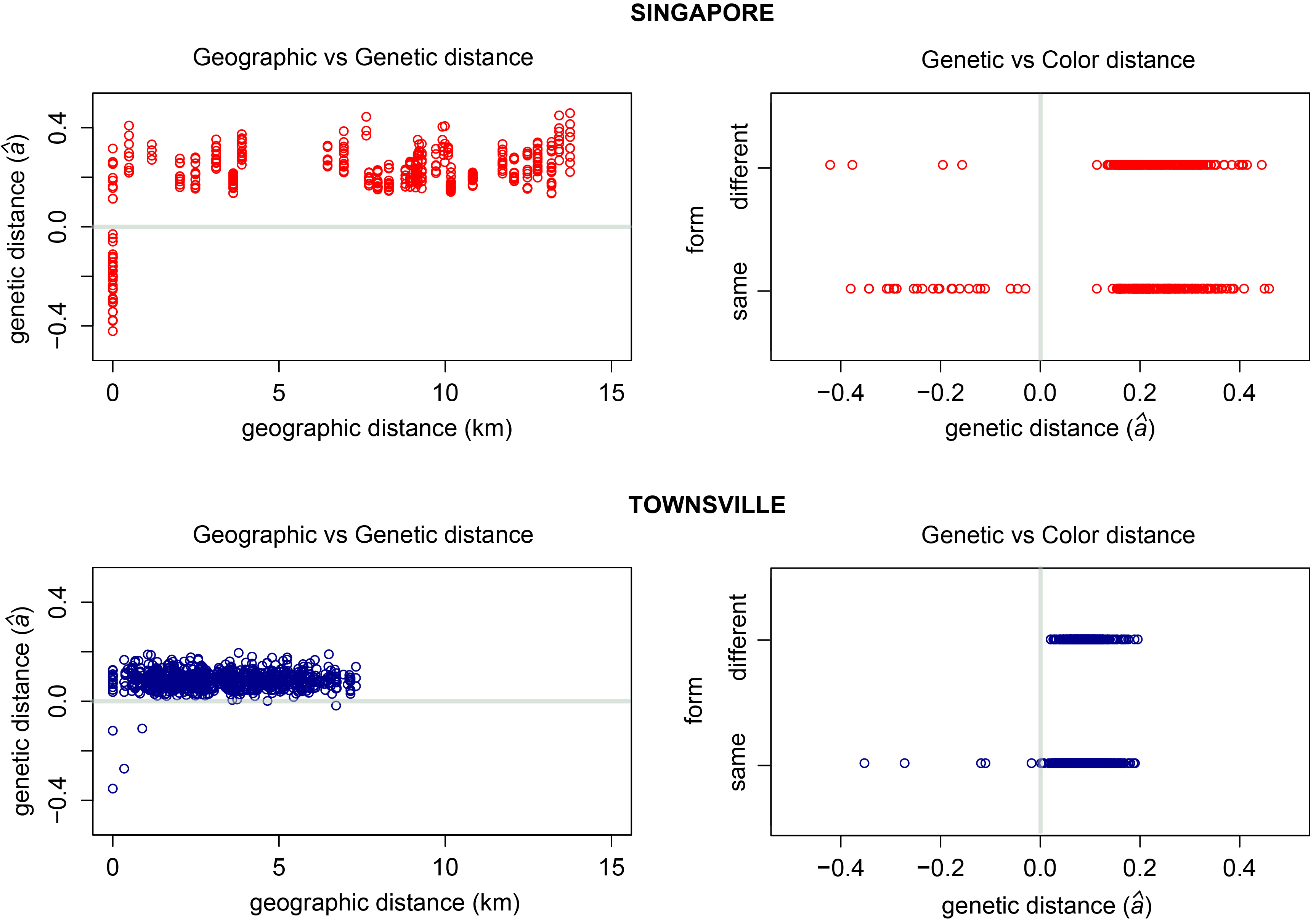
Pairwise genetic *versus* geographic and color distance. Pairs of *Aedes aegypti* collected in Singapore (upper graphs) and Townsville (lower graphs). A value of zero for Rousset’s genetic distance (*â*) indicates a distance between a pair of individuals randomly drawn from a given sample, while a negative value indicates lower than average genetic distance between a pair (i.e. their higher relatedness). Color distance between pairs of individuals was treated as a binary value: 0 (same color/form) and 1 (different color/form).

Individuals from northern Queensland were grouped into three clusters corresponding to the three towns where the sampling occurred (Figure 3c). A single exception was one *queenslandensis* individual from Cairns that was grouped with the *type* individuals from Gordonvale (Figure 3c). The two forms in Townsville could not be distinguished based on their nuclear genome-wide variation (Figure 3c). We found four pairs of closely related individuals: two *queenslandensis* and two *type* pairs (Figure 4, S4 file). In other words, all related pairs detected in Townsville were of the same form.

A lower frequency of related individuals in Townsville when compared to Singapore is not surprising given that different sampling methods were employed in these locations. Collection of multiple larvae from the same breeding container increases the chance of sampling family groups, as seen in Singapore and parts of Rio de Janeiro [21]. On the other hand, when BG-sentinel traps are used, the likelihood of related individuals being collected is low. In Townsville, 12.5% of pairs from the same trap were close relatives. Sampling effects are reflected in an elevated level of pairwise genetic distance over geographic distance for mosquitoes from Singapore when compared to Townsville (Figure 4). Such differences in genetic patterns could be erroneously interpreted as differences in the underlying processes (e.g. different dispersal rates), and highlight that sampling methods are crucial when inferring processes within and among *Ae. aegypti* populations.

In summary, we did not find any association between nuclear genetic variation and color/scaling variation in *Ae. aegypti* from Singapore and northern Queensland. Our results are unlikely to be caused by a lack of power to detect genetic structure, given that more than 16,000 genome-wide SNPs allowed us to delineate family groups at a very fine spatial scale. In fact, several families had the *queenslandensis* and *type* members. Given that a recent study of global *Ae. aegypti* populations at 12 microsatellite loci found that *Ae. aegypti formosus* and *Ae. aegypti aegypti* constitute one genetic cluster in Africa [37], it is not surprising that the two *Ae. aegypti aegypti* varieties also form one genetic cluster.

### *Wolbachia* infection

In addition to the absence of genetic structuring that corresponds to the two *Ae. aegypti aegypti* forms, another line of evidence in support of ongoing interbreeding is the presence of *Wolbachia* in both forms. We detected this endosymbiotic bacterium in three (out of four) *queenslandensis* individuals from Cairns and 14 (out of 15) *type* individuals from Gordonvale, using a light-cycler assay for *Wolbachia* detection [38]. *Wolbachia* is not naturally found in *Ae. aegypti*, but was introduced into the populations in Gordonvale in 2011 and Cairns in 2013 in an effort to reduce dengue transmission [39][40]. This was done by releasing *Wolbachia*-infected females and males from the colony that originated from the *type Ae. aegypti* [41]. Because the infection is transmitted from mother to offspring, the only way *queenslandensis* individuals could have become infected by *Wolbachia* is by mating with infected, *type* females. Given the high *Wolbachia* frequency (>85%) in Cairns and Gordonvale at the time of our sampling [42], finding the infection in 3/4 individuals caught in Cairns, and 14/15 individuals caught in Gordonvale was expected.

## Conclusion

Our analyses of mitochondrial and nuclear genome-wide variation and the *Wolbachia* infection indicate that *Ae. aegypti queenslandensis* and *Ae. aegypti type* mosquitoes interbreed freely, at least in Singapore and northern Queensland. These findings are of practical importance for control strategies that rely on successful mating between the released and target mosquitoes. Our results also re-enforce the recommendations by the early taxonomic authorities (Mattingly and McClelland) that *Ae. aegypti queenslandensis* should <u>not</u> be ranked as a subspecies.

## Acknowledgements

We thank Carly Herbertson, Melanie Commerford and other Eliminate Dengue field assistants for providing *Aedes aegupti* from Townsville and Cairns. We also thank the NeCTAR Cloud for providing computational resources. This work is funded by the National Health and Medical Research Council and the National Environment Agency, Singapore. The funders had no role in study design, data analyses, interpretation or manuscript writing.

## Author Contributions

GR: generated, analyzed, interpreted data, and wrote the manuscript with the input from AAH and IF; IF: generated and processed NGS data; AGC: done some field collections and molecular lab work; DS: noted reoccurring very pale form in QLD, identified and pre-processed samples from QLD; AC; SG L-P, HCT: identification and pre-processing of samples from Singapore; AAH: conceived the study. All authors have read and approved the final manuscript version.

## Supporting Information

**S1 Table. Sample information for additional *Aedes aegypti***. Mosquitoes from Rio de Janeiro (Brazil), Gordonvale (northern Queensland), Ho Chi Minh city (Vietnam), used in the DAPC analysis.

**S2 File. Mitochondrial haplotypes**. FASTA file with mitochondrial sequences from *Aedes aegypti* (Hap1-24), the Liverpool strain, and three outgroups used in the RAxML phylogenetic reconstruction. Mitochondrial haplotypes were generated by concatenating 90 bp RAD sequences from: ND2 (1-90 bp), COXI (91-270 bp), ATP6 (271-450 bp), ND5 (451-630 bp), ND4 (631-720 bp), ND6 (721-810 bp), cytB (811-1080 bp), ND1 (1081-1170 bp).

**S3 File. Pairwise nucleotide difference (*p*-distance) between mitochondrial haplotypes**. The number of base differences per site from between sequences are shown. The analysis involved 28 nucleotide sequences. All ambiguous positions were removed for each sequence pair. There were a total of 1170 positions in the final dataset. Evolutionary analyses were conducted in MEGA6.

**S4 File. Pairwise Relatedness**. Estimates of relatedness (*r*) by Wang (2002) for *Aedes aegypti* pairs in Singapore and Townsville. Reference: Wang, J. 2002. An estimator for pairwise relatedness using molecular markers. Genetics 160: 1203-1215.

**S1 Fig. Data quality parameters**. Boxplots of per-individual values for the proportion of uniquely aligned reads, RAD tag read depth, proportion of heterozygous loci, proportion of missing data for *Aedes aegypti* from Singapore (left) and Queensland (right).

## References

1. Kraemer MUG, Sinka ME, Duda KA, Mylne AQN, Shearer FM, Barker CM, et al. The global distribution of the arbovirus vectors Aedes aegypti and Ae. albopictus. Jit M, editor. Elife. eLife Sciences Publications, Ltd; 2015;4: e08347. doi:10.7554/eLife.08347

2. Benelli G, Mehlhorn H. Declining malaria, rising of dengue and Zika virus: insights for mosquito vector control. Parasitol Res. 2016;115: 1747–1754. doi:10.1007/s00436-016-4971-z

3. Halstead SB. Reappearance of Chikungunya, Formerly Called Dengue, in the Americas. Emerg Infect Dis. Centers for Disease Control and Prevention; 2015;21: 557–561. doi:10.3201/eid2104.141723

4. Chan M. Yellow fever: the resurgence of a forgotten disease. Lancet. Elsevier; 2016;387: 2165–2166. doi:10.1016/S0140-6736(16)30620-1

5. Mattingly PF, others. Genetical Aspects of the Aedes aegypti Problem. I.-Taxonomy and Bionomics. Ann trop Med Parasit. 1957;51: 392–408.

6. Howard LO, Dyar HG, Knab F. The Mosquitoes of North and Central America and the West Indies [Internet]. Issue 159,. Nature. Washington: Carnegie Institute; 1917. doi:10.1038/091420b0

7. McClelland GAH. A worldwide survey of variation in scale pattern of the abdominal tergum of Aedes aegypti (L.) (Diptera: Culicidae). Trans R Entomol Soc London. Blackwell Publishing Ltd; 1974;126: 239–259. doi:10.1111/j.1365-2311.1974.tb00853.x

8. Hill GF. Notes on Some Unusual Breeding-Places of Stegomyia Fasciata, Fabr., in Australia. Ann Trop Med Parasitol. Taylor & Francis; 1921;15: 91–92. doi:10.1080/00034983.1921.11684253

9. Powell JR, Tabachnick WJ. History of domestication and spread of Aedes aegypti––a review. Mem Inst Oswaldo Cruz. 2013;108 Suppl: 11–17. doi:10.1590/0074-0276130395

10. Moore M, Sylla M, Goss L, Burugu MW, Sang R, Kamau LW, et al. Dual African Origins of Global Aedes aegypti s.l. Populations Revealed by Mitochondrial DNA. PLoS Negl Trop Dis. 2013;7. doi:10.1371/journal.pntd.0002175

11. Brown JE, Evans BR, Zheng W, Obas V, Barrera-Martinez L, Egizi A, et al. Human impacts have shaped historical and recent evolution in Aedes aegypti, the dengue and yellow fever mosquito. Evolution. 2014;68: 514–525. doi:10.1111/evo.12281

12. Brown JE, McBride CS, Johnson P, Ritchie S, Paupy C, Bossin H, et al. Worldwide patterns of genetic differentiation imply multiple “domestications” of Aedes aegypti, a major vector of human diseases. Proc Biol Sci. 2011;278: 2446–2454. doi:10.1098/rspb.2010.2469

13. Chan A, Chiang L-P, Hapuarachchi HC, Tan C-H, Pang S-C, Lee R, et al. DNA barcoding: complementing morphological identification of mosquito species in Singapore. Parasit Vectors. London: BioMed Central; 2014;7: 569. doi:10.1186/s13071-014-0569-4

14. Wilson JJ. DNA Barcodes for Insects BT - DNA Barcodes: Methods and Protocols. In: Kress JW, Erickson LD, editors. Totowa, NJ: Humana Press; 2012. pp. 17–46. doi:10.1007/978-1-61779-591-6_3

15. Mayr E. Animal species and evolution. The Eugenics review. 1963. pp. 226–228. doi:10.1016/0169-5347(94)90187-2

16. Pennetier C, Warren B, Dabiré KR, Russell IJ, Gibson G. “Singing on the Wing” as a Mechanism for Species Recognition in the Malarial Mosquito Anopheles gambiae. Curr Biol. 2010;20: 131–136. doi:http://dx.doi.org/10.1016/j.cub.2009.11.040

17. Axford JK, Ross PA, Yeap HL, Callahan AG, Hoffmann AA. Fitness of wAlbB Wolbachia Infection in Aedes aegypti: Parameter Estimates in an Outcrossed Background and Potential for Population Invasion. Am J Trop Med Hyg. 2016;94: 507–516.

18. Chambers EW, Hapairai L, Peel BA, Bossin H, Dobson SL. Male Mating Competitiveness of a Wolbachia-Introgressed Aedes polynesiensis Strain under Semi-Field Conditions. Aksoy S, editor. PLoS Negl Trop Dis. San Francisco, USA: Public Library of Science; 2011;5: e1271. doi:10.1371/journal.pntd.0001271

19. Zhang D, Lees RS, Xi Z, Gilles JRL, Bourtzis K. Combining the Sterile Insect Technique with Wolbachia Based Approaches: II-A Safer Approach to Aedes albopictus Population Suppression Programmes, Designed to Minimize the Consequences of Inadvertent Female Release. PLoS One. Public Library of Science; 2015;10: e0135194. Available: http://dx.doi.org/10.1371%2Fjournal.pone.0135194

20. Massonnet-Bruneel B, Corre-Catelin N, Lacroix R, Lees RS, Hoang KP, Nimmo D, et al. Fitness of Transgenic Mosquito Aedes aegypti Males Carrying a Dominant Lethal Genetic System. PLoS One. Public Library of Science; 2013;8: e62711. Available: http://dx.doi.org/10.1371%2Fjournal.pone.0062711

21. Rasic G, Filipovic I, Weeks AR, Hoffmann A a. Genome-wide SNPs lead to strong signals of geographic structure and relatedness patterns in the major arbovirus vector, Aedes aegypti. BMC Genomics. 2014;15: 275. doi:10.1186/1471-2164-15-275

22. Behura S, Lobo N, Haas B. Complete sequences of mitochondria genomes of Aedes aegypti and Culex quinquefasciatus and comparative analysis of mitochondrial DNA. Insect Biochem …. 2011;41: 770–777. doi:10.1016/j.ibmb.2011.05.006

23. Behura SK, Lobo NF, Haas B, DeBruyn B, Lovin DD, Shumway MF, et al. Complete sequences of mitochondria genomes of Aedes aegypti and Culex quinquefasciatus and comparative analysis of mitochondrial dna fragments inserted in the nuclear genomes. Insect Biochem Mol Biol. 2011;41: 770–777. doi:10.1016/j.ibmb.2011.05.006

24. Langmead B, Trapnell C, Pop M, Salzberg S. Ultrafast and memory-efficient alignment of short DNA sequences to the human genome. Genome Biol. 2009;10. doi:10.1186/gb-2009-10-3-r25

25. Catchen J, Hohenlohe P a., Bassham S, Amores A, Cresko W a. Stacks: An analysis tool set for population genomics. Mol Ecol. 2013;22: 3124–3140. doi:10.1111/mec.12354

26. Rasic G, Schama R, Powell R, Maciel-de Freitas R, Endersby-Harshman NM, Filipovic I, et al. Contrasting genetic structure between mitochondrial and nuclear markers in the dengue fever mosquito from Rio de Janeiro: implications for vector control. Evol Appl. 2015;8: 901–915. doi:10.1111/eva.12301

27. Fontaine A, Filipovic I, Fansiri T, Hoffmann AA, Rasic G, Lambrechts L. Cryptic genetic differentiation of the sex-determining chromosome in the mosquito Aedes aegypti. bioRxiv. 2016; doi:http://dx.doi.org/10.1101/060061

28. Kalinowski ST. HP-RARE 1.0: A computer program for performing rarefaction on measures of allelic richness. Mol Ecol Notes. 2005;5: 187–189. doi:10.1111/j.1471-8286.2004.00845.x

29. Stamatakis A. RAxML version 8: A tool for phylogenetic analysis and post-analysis of large phylogenies. Bioinformatics. 2014;30: 1312–1313. doi:10.1093/bioinformatics/btu033

30. Hardy CM, Court LN, Morgan MJ, Webb CE. The complete mitochondrial DNA genomes for two lineages of Aedes notoscriptus (Diptera: Culicidae)itle. Mitochondrial DNA Part A DNA Mapping, Seq Anal. 2016;27: 2024–2025. doi:10.3109/19401736.2014.974171

31. Hardy CM, Court LN, Morgan MJ. The complete mitochondrial DNA genome of Aedes vigilax (Diptera: Culicidae). Mitochondrial DNA Part A DNA Mapping, Seq Anal. 2016;27: 2552–2553. doi:10.3109/19401736.2015.1038800

32. Jombart T, Devillard S, Balloux F. Discriminant analysis of principal components: a new method for the analysis of genetically structured populations. BMC Genet. BioMed Central Ltd; 2010;11: 94. doi:10.1186/1471-2156-11-94

33. Jombart T, Ahmed I. adegenet 1.3-1: new tools for the analysis of genome-wide SNP data. Bioinformatics. 2011;27: 3070–1. doi:10.1093/bioinformatics/btr521

34. Hardy OJ, Vekemans X. SPAGeDI: A versatile computer program to analyse spatial genetic structure at the individual or population levels. Mol Ecol Notes. 2002;2: 618–620. doi:10.1046/j.1471-8286.2002.00305.x

35. Moore M, Sylla M, Goss L, Burugu MW, Sang R, Kamau LW, et al. Dual African Origins of Global Aedes aegypti s.l. Populations Revealed by Mitochondrial DNA. PLoS Negl Trop Dis. 2013;7. doi:10.1371/journal.pntd.0002175

36. Mori A, Tsuda Y, Takagi M, Higa Y, Severson D. Multiple QTL Determine Dorsal Abdominal Scale Patterns in the Mosquito Aedes aegypti. J Hered. 2016;107: 438–444. doi:10.1093/jhered/esw027

37. Brown JE, Evans BR, Zheng W, Obas V, Barrera-Martinez L, Egizi A, et al. Human impacts have shaped historical and recent evolution in aedes aegypti, the dengue and yellow fever mosquito. Evolution (N Y). 2013;68: 514–525. doi:10.1111/evo.12281

38. Lee SF, White VL, Weeks AR, Hoffmann AA, Endersby NM. High-throughput PCR assays to monitor Wolbachia infection in the dengue mosquito (Aedes aegypti) and Drosophila simulans. Appl Environ Microbiol. 2012;78: 4740–4743. doi:10.1128/AEM.00069-12

39. Hoffmann a a, Montgomery BL, Popovici J, Iturbe-Ormaetxe I, Johnson PH, Muzzi F, et al. Successful establishment of Wolbachia in Aedes populations to suppress dengue transmission. Nature. 2011;476: 454–457. doi:10.1038/nature10356

40. Frentiu FD, Zakir T, Walker T, Popovici J, Pyke AT, van den Hurk A, et al. Limited Dengue Virus Replication in Field-Collected Aedes aegypti Mosquitoes Infected with Wolbachia. PLoS Negl Trop Dis. Public Library of Science; 2014;8: e2688.

41. Walker T, Johnson PH, Moreira L a, Iturbe-Ormaetxe I, Frentiu FD, McMeniman CJ, et al. The wMel Wolbachia strain blocks dengue and invades caged Aedes aegypti populations. Nature. 2011;476: 450–453. doi:10.1038/nature10355

42. Hoffmann AA, Iturbe-Ormaetxe I, Callahan AG, Phillips BL, Billington K, Axford JK, et al. Stability of the wMel Wolbachia Infection following Invasion into Aedes aegypti Populations. PLoS Negl Trop Dis. Public Library of Science; 2014;8: e3115. doi:10.1371/journal.pntd.0003115

